# *daf-42* is an evolutionarily young gene essential for dauer development in *Caenorhabditis elegans*

**DOI:** 10.1101/2023.04.24.538107

**Authors:** Daisy S. Lim, Jun Kim, Wonjoo Kim, Nari Kim, Sang-Hee Lee, Daehan Lee, Junho Lee

## Abstract

Under adverse environmental conditions, nematodes arrest into dauer, an alternative developmental stage for diapause. Dauer endures unfavorable environments and interacts with host animals to access favorable environments, thus playing a critical role in survival. Here, we report that in *Caenorhabditis elegans*, *daf-42* is essential for development into the dauer stage, as the null mutant of *daf-42* exhibited a “no viable dauer” phenotype in which no viable dauers were obtained in any dauer-inducing conditions. Long-term time lapse microscopy of synchonized larvae revealed that *daf-42* is involved in developmental changes from the pre-dauer L2d stage to the dauer stage. *daf-42* encodes large, disordered proteins of various sizes that are expressed in and secreted from the seam cells within a narrow time window shortly before the molt into dauer stage. Transcriptome analysis showed that the transcription of genes involved in larval physiology and dauer metabolism are highly affected by the *daf-4*2 mutation. Contrary to the notion that essential genes that control the life and death of an organism may well be conserved across diverse species, *daf-42* is an evolutionarily young gene conserved only in the *Caenorhabditis* genus. Our study shows that dauer formation is a vital process that is controlled not only by conserved genes but also by newly emerged genes, providing important insights into evolutionary mechanisms.

## Introduction

Animals adapt various strategies to survive an unpredictable, turbulent environment in the wild. One of these strategies is diapause, a form of dormancy in which the animal enters genetically programmed developmental arrest (Hand *et al*. 2016). Animals enter diapause in response to conditions that are indicative of environmental changes in the future. For example, insects enter diapause at various stages of life, such as embryo, pupa and adult, in response to changes in photoperiod and temperature (Diniz *et al*. 2017; Tougeron 2019). Some species of fish, such as killifish or kmar, enter embryonic arrest to survive dry seasons (Wourms 1972; Mesak *et al*. 2015).

The nematode *Caenorhabditis elegans* has an alternative non-feeding diapause stage known as dauer (Cassada and Russell 1975). Under favorable environments rich in nutrients*, C. elegans* quickly develops into reproducing adults through larval stages L1 to L4 in 3 days. However, young L1 *C. elegans* larvae can integrate information from the external environment and develop into the pre-dauer L2d stage and then enter dauer arrest under adverse conditions, such as high population density, high temperature or low food (Golden and Riddle 1984). Dauer worms are stress-resistant, specialized for long-term survival and can live for up to several months until they encounter a favorable environment and resume development into the adult stage and reproduce. The dauer stage is tightly linked to the “boom-and-bust” life cycle of *C. elegans*, in which the worms grow and reproduce with explosive speed in favorable environments and disperse when the resources are exhausted (Frezal and Felix 2015). At the “bust stage”, the young larvae develop into the dauer stage in to survive starvation and disperse into other locations, possibly by associating with carriers, such as slugs or isopods (Lee *et al*. 2011). Because *C. elegans* is mostly found in dauer stage in the wild, the dauer stage is thought to be essential for species survival. (Barriere and Felix 2005).

The dauer stage is not unique to *C. elegans*, but a widely conserved feature in nematodes, including parasitic nematodes. The infective juvenile (iL3) of parasitic nematodes is analogous to the dauer stage in that both are non-feeding, developmentally arrested third larval stages that resume development when the worms find suitable environmental conditions (Crook 2014). Parasitic nematodes invade or exit host animals at the iL3 stage, and the dauer stage may be an intermediate step for the evolution of parasitism in nematodes, because dauer and iL3 stages are associated with other animals for dispersal and survival (Hotez *et al*. 1993; Ahmed *et al*. 2013).

Dauer formation in *C. elegans* has been extensively studied as a genetic model of developmental plasticity. Conserved signaling factors, including the *daf-2*/insulin-like signaling and *daf-7*/TGF-β signaling pathways, relay information on the nutritional state and external environment and converge on steroid hormone signaling to regulate dauer development (Hu 2007; Fielenbach and Antebi 2008). Downregulation of these signaling components leads to the development of the diapause stage. More than 30 genes that have been identified from these studies regulate the developmental switch between adult or dauer stages, and mutants of these genes have altered developmental trajectories. However, downstream factors that directly participate in dauer development after the developmental decision have remained elusive.

Essential genes refer to genes that are necessary for survival of an organism or a cell. Since alterations in these genes cause lethality or other difficulties that severely hinder survival, essential genes are conventionally thought to be evolutionarily conserved factors that are present across diverse species. However, recent studies indicate that newly evolved, young genes also play vital biological roles (Chen *et al*. 2010; Ding *et al*. 2010; Charrier *et al*. 2012; Dietz *et al*. 2021). In this study, we discovered that *daf-42*, a previously uninvestigated genus-specific gene, is essential for dauer development in *C. elegans*. Null mutants of *daf-42* display lethality in a stage-specific manner at dauer entry. Genetic studies have revealed that DAF-42 acts downstream of the developmental decision process between diapause and reproductive stages and is critical in the short window of time immediately before molting into the dauer stage. We also investigate the phylogenetic conservation of this gene across nematode species.

## Materials and Methods

### Molecular Biology

Plasmids were generated and modified using classical restriction enzyme-based subcloning methods and Q5 site-directed mutagenesis (E0554/New England Biolabs), respectively. The description of plasmids and their generation is available in Supplementary Table 1.

### Worm maintenance and strains

*C. elegans* worms were grown in standard conditions (Brenner 1974), except for the strains in *daf-2(e1370)* and *daf-7(ok3125)* mutant backgrounds, which were raised at 15°C. The following strains were used:

Wild-type strain: N2, CB4856

Single mutant: CB1370 (*daf-2(e1370)*), LJ2701 (*daf-42(ys54)*), RB2302 (*daf-7(ok3125)*), DR2281 (*daf-9(m450)*)

Double mutant: LJ2703 (*daf-2(e1370); daf-42(ys54)*), LJ2770 (*daf-7(ok3125)*)*; daf-42(ys54),* LJ2771 (*daf-9(+/m450); daf-42(ys54)*), LJ2741 (*daf-2(e1370);daf-42(ys55)*), LJ2742 (*daf- 2(e1370);daf-42(ys58)*)

Transgenic strain:

Transgenic strains used in this study are listed in Supplementary Table 2.

### Generation of transgenic lines

Transgenes were introduced into worms by injecting purified plasmid DNA into the gonads of young adult hermaphrodite worms as described (Mello *et al*. 1991). Plasmid concentrations are listed in Supplementary Table 2.

### Dauer induction

N2 wild-type worms were raised in pheromone plates at 25°C to induce dauer stage. Approximately seven young adult worms were incubated at 25°C on pheromone plates seeded with OP50. Dauer worms on the plate were observed on day 5. The pheromone plates were prepared using synthetic dauer pheromones—ascaroside C7 (ascaroside 3, daumone 1), ascaroside C6 (ascaroside 1, daumone 2) and ascaroside C9 (ascaroside 2, daumone 3) (Jeong *et al*. 2005)— 10μM of which were added to growth media (NGM) without peptone.

Worms in *daf-2(e1370)*, *daf-7(ok3125)* and *daf-9(m540)* mutant backgrounds were raised in regular NGM plates seeded with OP50 at 25°C, as pheromone plates are unnecessary for these strains to induce the dauer stage.

### Transmission electron microscopy

*C. elegans* animals were overlaid with 20% bovine serum albumin in M9 buffer and frozen under high-pressure using a Leica EM HPM100 system (Leica, Austria). Animals were transferred to the freeze substitution apparatus (Leica EM AFS, Austria) in liquid nitrogen into a solution containing 2% osmium tetroxide and 1% water in acetone. The samples were maintained at – 90°C for 72 h, slowly warmed to –20°C (5 degree per hour), maintained for 24 h and slowly warmed to 0°C (6 degree per hour). The samples were washed three times with cold acetone at 0°C, transferred to room temperature, infiltrated with Embed-812 resin series by 20% increments in acetone (each step was 1 h) and embedded in Embed-812 (EMS, USA). After polymerization of the resin at 60°C for 36 h, sections were cut with a diamond knife on an ULTRACUT UC7 ultramicrotome (Leica, Austria) every 40 μm along the worm body. The 70-nm-thick sections were then mounted on formvar/carbon-coated grids, and were stained with 4% uranyl acetate for 10 min and lead citrate for 10 min. The samples were observed using a Tecnai G2 Spirit Twin transmission electron microscope (Thermo Fisher scientific, USA).

### Dead L2d development and temperature-shift assay

Synchronized embryos were obtained by placing day 2 adult worms (*n*=10–20) on a fresh NGM plate with OP50 for 2 hours at 15°C. The adults and the plates with newly laid embryos (eggs) were placed at 25°C for the desired time period to observe the developmental outcomes. L2d, dauer and dead L2d worms were distinguished based on their appearance and behavior, such as thin and long body, pointy head, lethargic behavior and rapid movement in response to touch using platinum pick. To calculate the ratio of each developmental stage, we divided the number of worms in each stage by the total number of worms visible on the plate. Some dauer worms were lost as they escaped the plate; these numbers were not included in the number of dauer worms or the total number of worms.

For the temperature-shift assay, synchronized embryos were obtained. Then, plates were kept at 15°C and moved to 25°C at desired time points. The developmental outcomes were observed at 120 hours after the eggs were laid (HAE).

### Microscopy

A stereo microscope (Zeiss Stemi 2000-C), fluorescence stereo microscope (Leica M205 FA) and confocal microscope (ZEISS LSM700) were used to observe *C. elegans*. Softwares LAS X (Leica) and Zen (black edition, Carl Zeiss) were used to obtain digital images using Leica M205 FA and Zeiss LSM700, respectively. For confocal imaging, worms were harvested with M9 buffer and placed on 3% agar pads with 3mM levamisole.

### Mutation mapping through mapping-by-sequencing

To identify the causative mutation for the lethal phenotype, we followed the mapping-by-sequencing strategy (Doitsidou *et al*. 2010; Doitsidou *et al*. 2016). *daf-2(e1370); daf-42(ys54)* mutant was mated with CB4856 to obtain 48 F2 hybrid strains that developed into dead L2d at 25°C. The worm strains were pooled, and genomic DNA was extracted with a Qiagen Gentra Puregene Tissue Kit using “Purification of DNA from nematodes” protocol. Macrogen Inc. (South Korea) prepared the library and sequenced the pooled genomic DNA using the Illumina Hiseq 4000 100PE platform. Approximately 5 Gb of sequencing data were obtained.

Using the whole-genome sequencing data, the approximate location of the *ys54* mutation was identified using the MimodD pipeline (http://mimodd.readthedocs.io/en/latest/index.html, DOI: <u>10.5281/zenodo.1189838</u>). Using this pipeline, WGS reads were aligned to the WS220 reference genome, and the density of single nucleotide polymorphisms (SNPs) from CB4856 were calculated and visualized. A list of mutations that could be *ys54* was obtained from the 5 Mb to 11 Mb region of chromosome IV, which showed an extremely low density of SNPs from CB4856.

### Search for secreted, large and disordered proteins

Data on *C. elegans* proteins, including signal peptide prediction, length and disorder content were obtained from MobiDB (https://mobidb.bio.unipd.it). Proteins that have predicted signal peptides, length above 1,000 residues and over 50% disorder content predicted by MobiDB-lite were filtered. MobiDB-lite, version 3.10.0, was used to predict positions of disordered regions in DAF-42 homologs.

### Generation of new alleles of *daf-*42 using CRISPR/Cas9

A co-conversion strategy using CRISPR/Cas9 was employed to generate new *daf-42* mutants (Arribere *et al*. 2014; Ward 2015). 40 ng/μL of each plasmids containing CRISPR/Cas9 and sgRNA for *daf-42*—pJL1821. pJL1823, pJL1824—were injected into gonads of N2 worms with 20 ng/ul of pJA42 and 20 ng/μL of AF-JA-53 (injection marker containing *rol-6(su1006)* and repair template to generate *rol-6(su1006)*, respectively) (Arribere *et al*. 2014). Injected worms were raised at 25°C, and the F1 offsprings with roller phenotype were singled out for propagation and checked for mutations at *daf-42* locus using T7E1 assay to select heterozygote mutants. Then, non-roller F2 offsprings of the selected heterozygote F1 worms were screened for homozygous mutation at *daf-42* using T7E1 assay.

### Scoring of transgenic animals

Synchronized larvae were obtained by placing ten day 2 adult worms on a fresh NGM plate at 15°C for 2 hours and then incubated at 25°C for dauer induction. To observe phenotypes of transgenic animals, transgenic L2d worms were first selected at 48 HAE under a fluorescence stereo microscope (Leica M205 FA). The selected worms were raised in NGM plates with OP50 for desired times until they were observed for scoring of the dead L2d phenotype.

To score expression status, the transgenic worms that express *daf-42p::signal peptide::mCherry* and *grd-10p::gfp* transgenes were observed under a fluorescence stereo microscope (Leica M205 FA). At 48 HAE, each transgenic worm was transferred to a fresh, labeled NGM plate with OP50 for observation. The expression pattern of *daf-42p::signal peptide::mCherry* was observed for each worm every four hours until 72 HAE. Dauer worms tend to move out of the *E. coli* food lawn, and sometimes escape the agar NGM media. Cases in which dauer larvae were not found, were recorded as “worm not found”.

### RNA sequencing and analysis

Synchronized *daf-2* and *daf-2; daf-42* embryos were obtained by placing 20 adult worms each on six NGM plates with OP50 for 2 hours at 15°C and then incubated at 25°C for dauer induction.

At 52 HAE and 60 HAE, the worms were harvested with M9 and washed with distilled water five times. The, 250 μL of TRIzol was added to 50 μL of washed worm harvest, and the worm samples were freeze-thawed 10–15 times for disruption. Harvest *daf-2; daf-42* at 60 HAE included both live and dead L2d worms. RNA was extracted through chloroform—isopropanol precipitation. To minimize variation, each set of *daf-2* and *daf-2; daf-42* at 52 and 60 HAE was grown, harvested and disrupted simultaneously. Triplicates of each set were made.

Theragen Bio Inc. (South Korea) prepared a library with TruSeq Stranded mRNA Sample Prep Kit and RNA sequencing on the Illumina NovaSeq 6000 150-bp paired-end platform. A total of 63–90 million reads were obtained from each sample. The RNA-seq data were analyzed using Kallisto and Sleuth (Bray *et al*. 2016; Pimentel *et al*. 2017).

### Protein alignment

Protein sequences from 202 bioprojects of 163 nematodes and platyhelminthes were obtained from WormBase Parasite 16 (https://parasite.wormbase.org/ftp.html) (Howe *et al*. 2015; Howe *et al*. 2017) and aligned with the DAF-42 m protein sequence using DIAMOND in the ultra- sensitive mode (Buchfink *et al*. 2021). For every species, one protein with the highest bit score was selected as the *daf-42* homolog.

To analyze if a homolog has two sites of alignment at the N- and C-terminus, two overlapping fragments of DAF-42 m (1–1500 aa and 1001–2402 amino acid residues) were aligned to protein sequence of *daf-42* homologs in *Caenorhabditis* species, under same procedure with DIAMOND.

Protein dot plot was generated using EMBOSS Dotmatcher (https://www.ebi.ac.uk/Tools/seqstats/emboss_dotmatcher/), under window size 10, threshold 23 and matrix BLOSUM62 (Madeira *et al*. 2022). Protein sequences of *C. elegans* DAF-42 m and its best-matching homologs in *Caenorhabditis* species were used as input.

## Results

### Identification of a mutant with stage-specific lethal phenotype during dauer entry

In harsh environments, young *C. elegans* L1 larvae choose to develop into pre-dauer L2d and then arrest into dauer stage until conditions become favorable for reproduction (Fig. 1a). Interestingly, during a mutant screen for dauer-stage behavior (Lee *et al*. 2017a), we discovered that the mutant strain RB2489 developed normally into reproducing adults but failed to develop into the dauer stage and showed a “no viable dauer” phenotype. Larvae grown in a dauer-forming condition (high concentration of dauer pheromone, high temperature and little food) developed into corpses (Supplementary Fig. 1, a and b). These dead worms did not resume growth after transfer into the adult-inducing normal condition, even after one or two days. The dead worms were immobile and trapped in their own cuticles in a straight or slightly curved posture, unlike the S-shaped wavy posture of live worms. This completely penetrant, stage-specific lethal phenotype was unlike the dauer-constitutive (Daf-c) and dauer-defective (Daf-d) phenotypes of known mutants of dauer formation (*daf)* genes, which control the developmental decisions between diapause and reproductive growth; *daf-c* and *daf-d* mutant larvae develop into dauer arrest in absence of pheromones and into reproductive stages even in the presence of pheromones, respectively (Riddle *et al*. 1981; Fielenbach and Antebi 2008). Outcrossing with wild-type N2 revealed that the phenotype was independent of the *ins-15(ok3444)* mutation of strain RB2459 (Consortium 2012). Therefore, we named the causative gene and mutation *daf-42(ys54)*.

**Figure 1.**
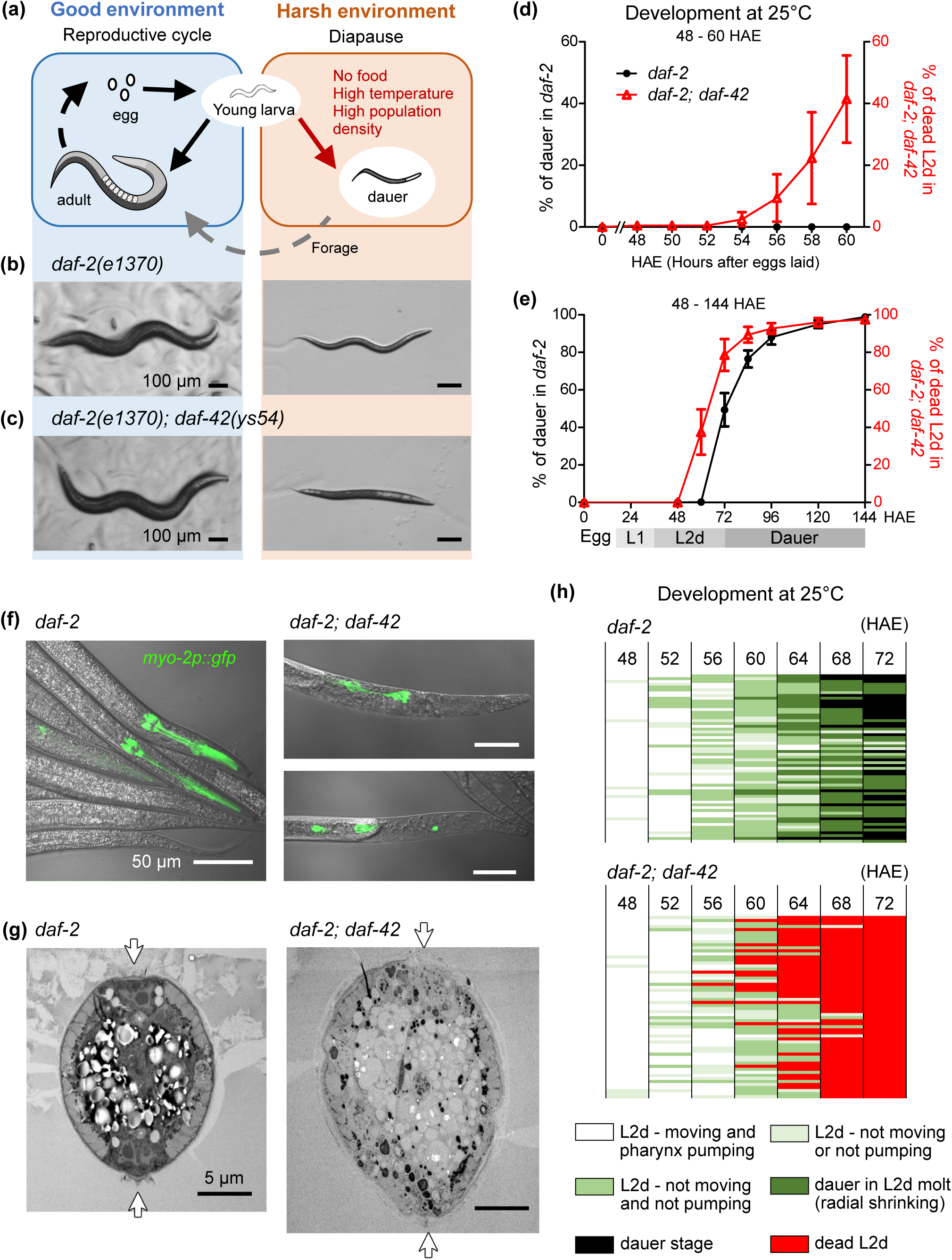
The *daf-42(ys54)* mutant displays a lethal phenotype uner dauer-inducing conditions. (a) Life cycle of *Caenorhabditis elegans*. Young larvae in harsh environments develop into L2d and then to dauer larvae, instead of quick growth into reproductive adults in favorable environments. (b) *daf-2(e1370)* worms develop into reproductive adults at 15°C (left) and into dauer larvae at 25°C (right). (c) *daf-2(e1370); daf-42(ys54)* worms develop into adults at 15°C (left) but fail to develop into dauer stage at 25°C (right). (d, e) Percent of worms that develop into dauer stage in *daf-2* worms (black) and that fail to develop into the dauer stage in *daf-2; daf-42(ys54)* worms (red) at 25°C during 48 to 144 hours after the eggs were laid (HAE) (d) and 48 to 60 HAE (e). n > 350 for (d) and > 560 for (e) in three trials. Data show mean ± SEM. (f) Transgenic *daf-2(e1370)* worms (left) and *daf-2(e1370); daf-42(ys54)* worms (right above, right below) expressing gfp in the pharyngeal muscle (*myo-2p::gfp*) when grown at 25°C for 72 HAE. (g) Electron microscopy images of cross sections of *daf-2(e1370)* worms (left) and *daf- 2(e1370);daf-42(ys54)* worms (right) in the mid-body when grown at 25°C for 96 HAE. White arrows indicate positions of dauer alae, which has formed in *daf-2(e1370)* control (left), but not in *daf-2(e1370); daf-42(ys54)* mutant (right). (h) Development of *daf-2(e1370)* (above) and *daf-2(e1370); daf-2(ys54)* (below) at 25°C from 48 to 72 HAE. Each row represents a single worm. After the lethargus period (light green), *daf- 2(e1370)* larvae undergo radial constriction of the body and ecdysis (dark green) followed by complete maturation into the dauer stage (black). However, *daf-2(e1370); daf-42(ys54)* larvae show developmental defects (red) after the lethargus period. n = 60 in three trials

The *daf-42(ys54)* mutation also caused the same phenotype in the *daf-2(e1370)* mutant background (Fig. 1, b and c), a temperature-sensitive mutant that constitutively develops into the dauer stage in high temperature (25°C) even in the presence of food and absence of dauer pheromones (Gottlieb and Ruvkun 1994). This indicates that in addition to changes in rearing conditions, the “no viable dauer” phenotype can be elicited by genetic manipulations that induce development into the dauer stage. We continued to study *daf-42* in the *daf-2(e1370)* mutant background as it was easier to induce dauer stage and the induced dauer larvae were relatively more uniform in size and development than the larvae with the wild-type background.

To investigate the mutant phenotype, we collected synchronized embryos and observed their development at 25 °C and various time points. *daf-2* control worms developed into the dauer stage after 60 hours after eggs were laid (HAE), and approximately one-half of the population developed into dauer by 72 HAE. The *daf-2; daf-42* mutant worms started developing the defective phenotype at 54 HAE, and nearly 80% of the population developed the phenotype by 72 HAE, which correspond to the end of the L2d stage (Fig. 1, d and e, Supplementary Fig. 2, a and b). Overall, the developmental defect phenotype of *daf-2; daf-42* mutant larvae appears 8– 12 hours prior to the completion of dauer development in control *daf-2* worms.

The *daf-2; daf-42* mutant was no different from the *daf-2* control at 48 HAE; however, at 72 HAE and 96 HAE, *daf-2; daf-42* mutant worms had developmental defects, such as degraded pharyngeal muscle and shortened head (Fig. 1f and Supplementary Fig 2, c-f). Although the worms were alive and could move their heads within the cuticle at this point, they were unable to molt out of the old cuticle and develop into the dauer stage afterwards. Electron microscopy analysis of worm cross sections revealed that at 72 HAE, *daf-2; daf-42* mutant larvae had damage not only around the head, but also at the mid-body and tissues were loose, unlike the packed organization in *daf-2* (Supplementary Fig. 3). At 96 HAE, the *daf-2* control worm fully developed into dauer stage, showing dauer cuticle, alae and constricted body size (Wolkow and Hall 2011); however, the *daf-2; daf-42* mutant had damaged head region, ill-developed body and has not escaped the L2d cuticle (Fig. 1g, Supplementary Fig. 3). Moreover, in *daf-2; daf-42* mutant, dauer alae were not formed, and the striated layer in the dauer cuticle terminated at the seam cell area. These results indicate that *daf-42(ys54)* mutation causes various developmental defects during the L2d-to-dauer transition and that *daf-42* is essential gene during L2d for development into the dauer stage.

To identify when the developmental defect phenotype occurs, we examined the development of individual worms during the L2d-to-dauer transition. In *C. elegans*, molting can be separated into the lethargus phase and ecdysis. In the lethargus phase, worms slow down physical movement and become lethargic. In ecdysis, the worms resume physical movement and remove their old cuticle (Lazetic and Fay 2017). This pattern was visible in *daf-2* control worms, which showed 4–12 hours of lethargus period followed by ecdysis and maturation into the dauer stage (Fig. 1h, top, Supplementary Fig. 4). During ecdysis, we could see a thinner worm inside the L2d cuticle, which indicates that the worm went through radial body constriction at this point (Supplementary Fig. 4c). In contrast, *daf-2; daf-42* mutant worms do enter the lethargus phase but does not show radial constriction and ecdysis. Instead, the larvae develop the developmental defect phenotype (Fig. 1h, bottom, Supplementary Fig. 4g). This indicates that the *daf-42* mutant enters the L2d-to-dauer transition but is unable to develop into the dauer stage. Together, these results show that the *daf-42* mutant shows a developmental defect phenotype specifically at the L2d-to-dauer transition, which leads to lethality under dauer-forming conditions.

### *daf-42* acts in dauer development after developmental commitment to diapause

Various genetic pathways regulate dauer formation. We have previously shown that *daf-2; daf-42* double mutant worms exhibit a “no viable dauer” phenotype. Moreover, we analyzed the *daf-42* mutant phenotype in the context of *daf-7* and *daf-9*, which are upstream regulators of tgf-ß and nuclear hormone signaling pathways, respectively (Ren *et al*. 1996; Gerisch and Antebi 2004). *daf-7(ok3125)* is a temperature-sensitive daf-c mutant that constitutively develops into a dauer at 25°C. We found that the *daf-7(ok3125); daf-42(ys54)* double mutant exhibits the lethal phenotype at 25°C (Fig. 2, a and b). *daf-9(m540)* causes dauer formation unconditionally, followed by dauer exit and growth into reproductive adults after 1–2 days (Jia *et al*. 2002). We were unable to secure a *daf-42(ys54); daf-9(m540)* double mutant strain because worms committed to forming dauer became lethal (Fig. 2c). Further examination of 1,210 offspring from a *daf-42(ys54); daf-9(+/m540)* heterozygote mutant revealed 299 worms with the lethal phenotype, which follows the Mendelian inheritance and confirms that the *daf-42(ys54); daf- 9(m540)* double mutant also exhibits the “no viable dauer” phenotype. Our results indicate that *daf-42* functions after the decision process, which is regulated by dauer formation genes *daf-2*, *daf-7* and *daf-9*, and suggest that *daf-42* is a distinct type of daf gene that functions in dauer development during the L2d-to-dauer transition after the decision to enter the dauer stage has been made.

**Figure 2.**
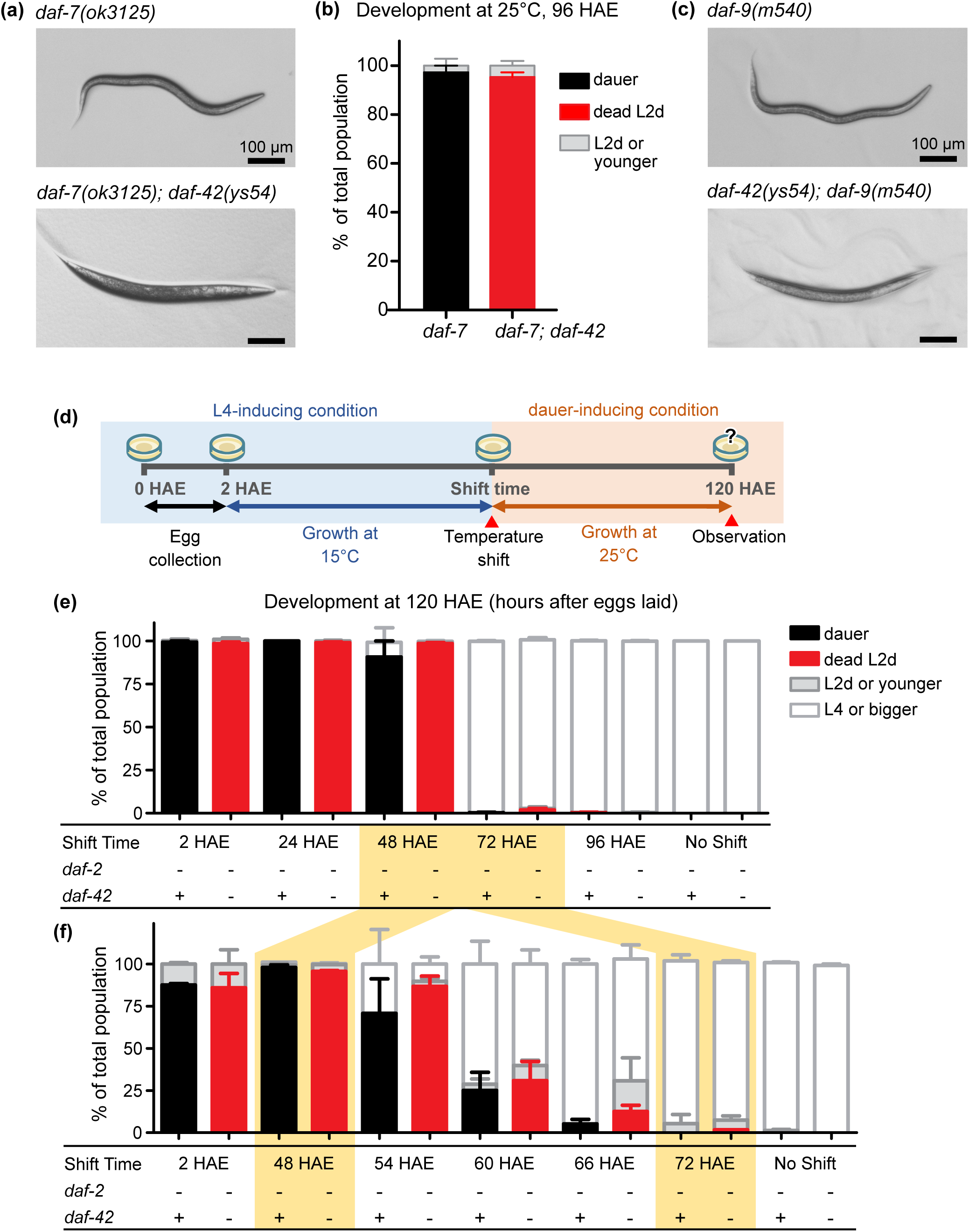
*daf-42* functions downstream of dauer commitment. (a) Representative images of the development of *daf-7(ok3125)* into dauer stage (above) and *daf- 7(ok3125); daf-42(ys54)* into dead L2d (below) at 25°C at 96 HAE. (b) The *daf-7(ok3125)* mutant develops into the dauer stage at 25°C, whereas a similar percentage of the *daf-7(ok3125); daf-42(ys54)* double mutant develops into dead L2d in the same condition. Data show mean ± SEM. n > 118 in three trials. (c) Representative images of the development of *daf-9(m540)* into the dauer stage (above) and *daf-42(ys54); daf-9(m540)* into dead L2d (below) at 25°C at 96 HAE. (d) Schematic representation of the temperature-shift assay. Eggs were collected for 2 hours and grown at 15°C until they were transferred to 25°C at certain time points (shift time). The results on how the worms grew were taken at 120 HAE. (e, f) Temperature shift assays from L4-inducing condition (15°C) to dauer-inducing condition (25°C) for *daf-2(e1370)* and *daf-2(e1370); daf-42(ys54)* with shift times every 24 hours between 2 HAE and 120 HAE (e) and every 6 hours between 48 and 72 HAE (f). Similar percentages of dauer development in *daf-2* and dead L2d formation in *daf-2; daf-42* show that the lethal phenotype at dauer entry caused by the *daf-42* mutation is downstream of commitment into the dauer stage development. (+) indicates wild-type allele and (-) indicates mutant allele. Data show mean ± SEM. n > 500 for (e) and > 130 for (f) in three trials.

The decision to develop into pre-dauer L2d is made at L1 if the worm is in unfavorable condition. In other words, exposure to favorable conditions before late L1 can still induce diapause, on the condition that the larva is in an unfavorable condition by late L1 (Cassada and Russell 1975; Golden and Riddle 1984; Schaedel *et al*. 2012). To investigate whether the lethal phenotype is affected by the experience of favorable conditions, we performed a shift-to- unfavorable assay by shifting the growth temperature of synchronized embryos from 15°C to 25°C progressively (Fig. 2d). *daf-2* control worms transferred to 25°C after 54-72 HAE started to be unresponsive to 25°C and developed into reproductive stages, indicating developmental decision at late L1 (Fig. 2, e and f, black bars).

We hypothesized that if the “no viable dauer” phenotype occurs specifically the L2d-to-dauer transition after the decision into diapause, then the proportion of dauer worms in *daf-2* will be close to that of dead L2d worms in *daf-2; daf-42.* Similar to our hypothesis, the proportion of larvae entering reproductive fate and diapause fate in *daf-2; daf-42* matched that of *daf-2*, except that *daf-2; daf-42* mutants became dead L2d worms instead of dauer worms (Fig. 2, e and f, red bars). This shows that the dead L2d phenotype is unaffected by exposure to favorable environment before late L1 and suggests that *daf-42* facilitates physiological development that starts after the developmental decision to enter dauer during L2d.

### *daf-42* is a previously uninvestigated gene that encodes large, disordered proteins

To identify the causative mutation, we employed mapping-by-sequencing—a method that combines single nucleotide polymorphism (SNP) mapping with whole-genome sequencing (Doitsidou *et al*. 2010; Doitsidou *et al*. 2016). Mapping-by-sequencing followed by extensive outcrossing revealed a nonsense mutation in exon 1 of ORF Y40C5A.3, located in the middle of chromosome IV, as *daf-42(ys54)* (Fig. 3a, Supplementary Fig. 5, a and b, and Supplementary Table 3). Null mutations introduced into exon 1 of *daf-42* using CRISPR/Cas9, *ys55* and *ys58*, reproduced the lethal phenotype of *daf-42(ys54)*, suggesting that the disruption *daf-42* is responsible for the “no viable dauer” phenotype (Fig. 3, a–c). This was further confirmed by the complete rescue of the lethal phenotype in transgenic *daf-2; daf-42* mutants carrying a fosmid from the *C. elegans* genomic DNA library or a newly constructed plasmid containing the wild-type allele of this gene (Fig. 3, d and e). Thus, ORF Y40C5A.3 is *daf-42*.

**Figure 3.**
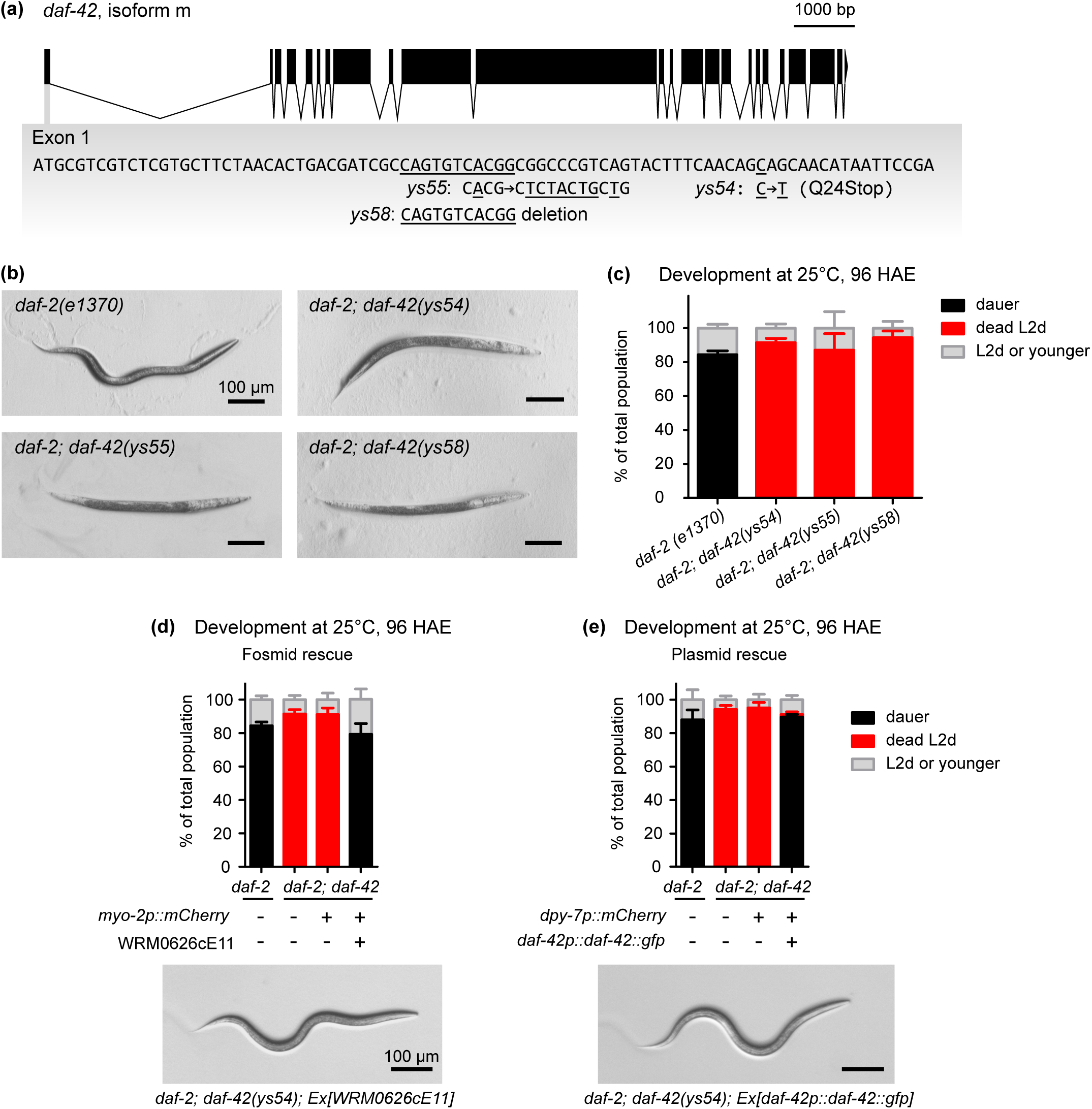
*ys54* is a nonsense mutation in *daf-42*—a previously uninvestigated gene in the middle of Chromosome IV. (a) Gene structure of *daf-42 m*, the isoform of *daf-42* that contains all exons. Black blocks indicate exons, and black lines indicate introns. Mutant alleles *ys54, ys55, ys58*, which are on exon 1, are indicated below. (b) Representative microscopy images of dauer development in *daf-2(e1370)* (left above), *daf- 2(e1370); daf-42(ys54)* (right above), *daf-2(e1370); daf-42(ys55)* (left below) and *daf-2(e1370); daf-42(ys58)* (right below). (c) Development of *daf-2(e1370)* control worms and *daf-2(e1370); daf-42* mutant worms at 25°C. *daf-2* develops into the dauer stage; however, all three null alleles of *daf-42* (*ys54*, *ys55* and *ys58*) cause development into dead L2d larvae. Data show mean ± SEM. n > 70 in three trials. (d, e) “No viable dauer” phenotype of *daf-2; daf-42(ys54)* larvae is rescued by transgenic fosmid WRM0626cE11 (d) and plasmid (e) that contain the gene *daf-42*. Representative pictures of the worm with rescued phenotype are shown in the picture below the graph. (+) indicates presence and (-) indicates the absence of each transgene. Data show mean ± SEM. n > 68 in three trials.

To our knowledge, *daf-42* has not been investigated to date. *daf-42* encodes large proteins, ranging from 1,101 to 2,402 amino acid residues across 17 isoforms (Supplementary Fig. 5c) (Harris *et al*. 2020). The largest isoform DAF-42 m contains all exons and is the 269th- largest protein among 26,548 proteins in the *C. elegans* proteome (Okimoto *et al*. 1992; Consortium 1998). Structural analyses, including the analysis by AlphaFold 2, indicated that DAF-42 m consists primarily of unstructured regions (Supplementary Fig. 5d) (Necci *et al*. 2017; Jumper *et al*. 2021; Piovesan *et al*. 2021). The first 15 amino acid residues were predicted to be a signal peptide, suggesting that the protein is not likely cytosolic (Supplementary Fig. 5d). To predict the functions of *daf-42,* we searched for large proteins that are predicted to be disordered and have a signal peptide. Interestingly, only seven proteins in *C. elegans* had these characteristics, four of which are incorporated into the cuticle or the extracellular matrix (Supplementary Table 4). This implies that *daf-42* may also play similar structural roles in these tissues.

### *daf-42* encodes secreted proteins that are essential during a narrow time window during dauer entry

To identify the role of *daf-42* in dauer development, we examined the site of *daf-42* expression. Interestingly, studies on the *C. elegans* transcriptome suggested that *daf-42* is specifically expressed during dauer entry and is not expressed at other developmental stages (Fig. 4a) (Lee *et al*. 2017b; Harris *et al*. 2020). Consistently, transgenic worms expressing mCherry under the *daf-42* promoter showed expression in seam cells between 48 and 60 HAE (Fig. 4b), which corresponds to the time of appearance of the dead L2d phenotype (Fig. 1d). Seam cells are specialized syncytial hypodermic cells along the length of the animal that synthesize cuticle components, such as collagen, and are important for epidermal elongation and molting (Singh and Sulston 1978; Thein *et al*. 2003). The expression of wild-type *daf-42* in seam cells partially rescued the phenotype of the *daf-2; daf-42* mutant, indicating that expression of *daf-42* in seam cells is essential for dauer development (Fig. 4c, Supplementary Fig. 6). The partial rescue of the phenotype using a constitutive seam cell promoter may suggest that expression in a narrow time window is important.

**Figure 4.**
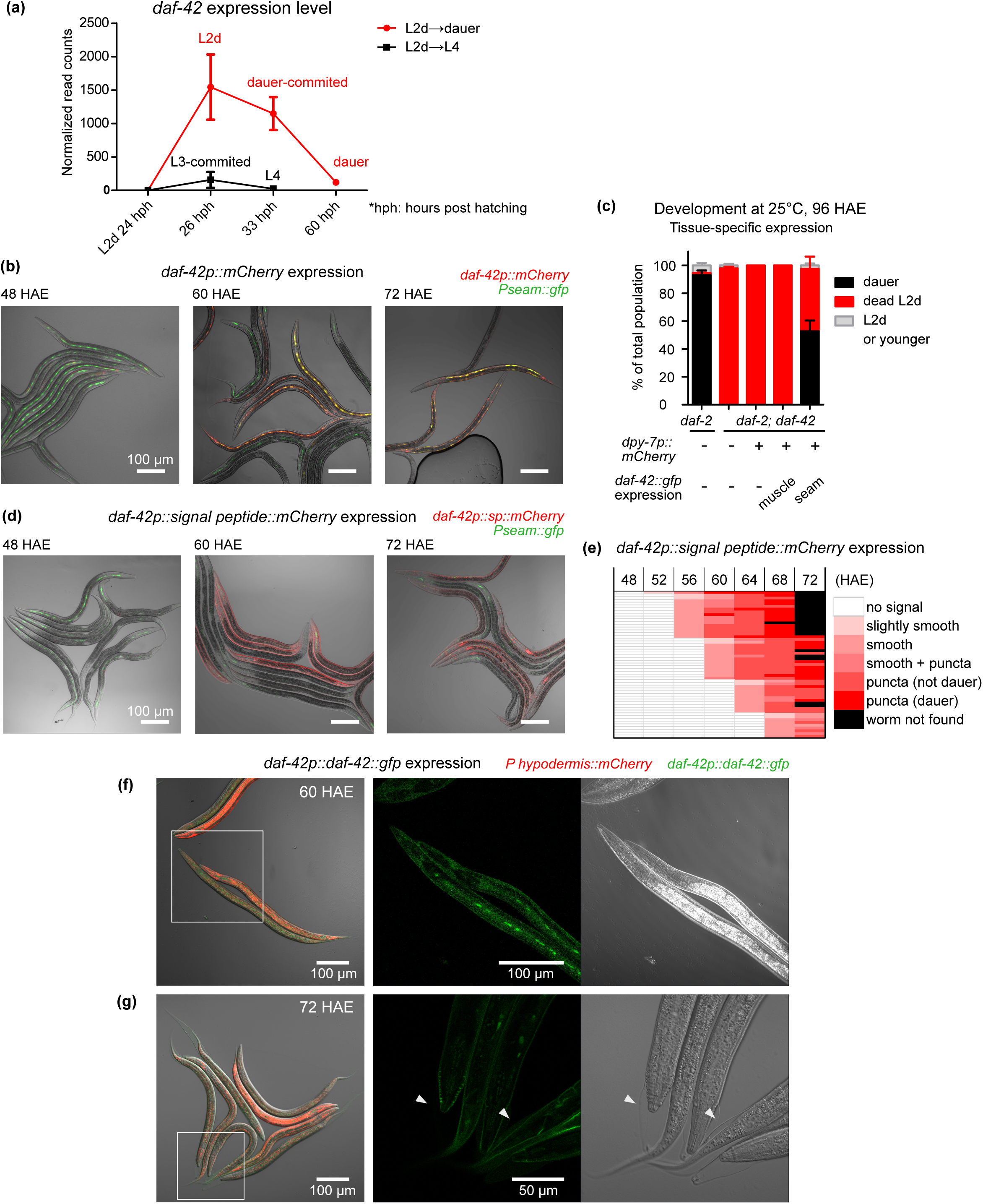
DAF-42 is secreted from hypodermis and acts in a narrow time window during L2d-to-dauer transition. (a) Expression level of *daf-42* in dauer-entering larvae and L4-developing larvae. *daf-42* is expressed specifically at dauer entry. Transcriptome data is from Lee et. al. (2017). Data show the mean ± SEM of the triplicates. (b) Representative images of *daf-2* expressing *daf-42p::mCherry* at 48 HAE (left), 60 HAE (middle) and 72 HAE (right) under dauer-inducing conditions (25°C). Expression pattern of *daf- 42p::mCherry* colocalize with seam cell marker *grd-10p::gfp*. (c) Ectopic expression of wild-type *daf-42* under the seam cell promoter *grd-10p* partially rescues the dead L2d phenotype of the *daf-42(ys54)* mutant. In contrast, the expression of the same transgene in body wall muscles using the *myo-3* promoter did not rescue the lethal phenotype. (+) indicates presence and (–) indicates absence of each transgene. Data show mean ± SEM. n > 80 in three trials. (d) Expression of mCherry under *daf-42* promoter and signal peptide (*daf-42p::signal peptide::mCherry*) in *daf-2(e1370)* background at 48 HAE (left), 60 HAE (middle) and 72 HAE (right) under dauer-inducing conditions (25°C). (e) Quantification of *daf-42p::sp::mCherry* expression between 48 and 60 HAE. Expression occurs between 52 and 64 HAE. Each row indicates a single larva. n = 53 in 3 trials. (f) Expression of *daf-42p::daf-42::gfp* in *daf-2(e1370)* larvae shows localization of DAF-42 in seam cells, hypodermis and worm surface at 60 HAE (f) and 72 HAE (g). White box in the left indicates the region with the enlarged image shown in the right. White arrowheads indicate places where detachment of dauer-forming worms from old L2d cuticles can be seen at 72 HAE.

We generated transgenic worms carrying the putative signal peptide fused to mCherry under the *daf-42* promoter (*daf-42p::signal peptide::mCherry*) to investigate the functionality of the predicted signal peptide (Supplementary Fig. 7a). Addition of a signal peptide to the N-terminus of mCherry drove its expression at the periphery of the worm body at 60 HAE, in stark contrast to the previous reporter assay without the signal peptide (Fig. 4d, Supplementary Fig. 7b). The mCherry signal was slightly visible in coelomocytes in some worms at 48 HAE, was more visible in the worm periphery at 60 HAE, and formed puncta through the worm body at 72 HAE as the worms developed into the dauer stage. Analysis of mCherry expression every 4 hours between 48 and 72 HAE revealed that in all cases, smooth expression in the periphery preceded puncta formation (Fig. 4e, Supplementary Fig. 7c). These results suggest that DAF-42 is secreted from seam cells to peripheral body at late L2d before before entering the dauer stage.

Interestingly, the time of expression of *daf-42p::signal peptide::mCherry* coincided with the appearance of the developmental defect in *daf-2; daf-42* mutants. For example, *daf-2; daf-42* mutant larvae started to exhibit the phenotype at 54 HAE (Fig. 1d); similarly, transgenic *daf-2* worms started to express mCherry as early as 52 HAE (Fig. 4e). 31.5% of transgenic *daf-42* worms had *daf-42* promoter activity at 56 HAE, and a similar percentage of *daf-2; daf-42* mutant worms had the developmental defect phenotype at 58 and 60 HAE (Fig. 1e, Fig. 4e). Thus, the absence of *daf-42* in a narrow time window during dauer entry results in developmental defects and suggests that *daf-42* plays a crucial role in dauer development shortly after its expression.

We analyzed localization of DAF-42 protein using a translational reporter line expressing *daf-42* fused to gfp (*daf-42::gfp)*. At 60 HAE, we observed expression in the seam cell, hypodermis and at the surface of the worm (Fig. 4f). At 72 HAE, ecdysis had begun, some worms had detached their head or tail from the old L2d cuticle (Fig. 4g, arrowheads), and the DAF-42::GFP signal did not remain in the L2d cuticle but was localized with the dauer worm inside the L2d cuticle (Fig. 4g). Together, our results show that *daf-42* is expressed during the L2d-to-dauer transition in seam cells and is secreted towards the surface of the dauer worm, such as the hypodermis or cuticle.

### Absence of *daf-42* interferes with transcriptomic changes during dauer entry

To examine the transcriptional changes accompanied by the absence of DAF-42 during dauer entry, we performed RNA-sequencing on *daf-2* and *daf-2; daf-42* worms at 52 and 60 HAE, the time at which 0% and ∼40% of *daf-2; daf-42* mutants form dead L2d at 25°C, respectively (Fig. 1e). Although the use of the *daf-2* background have unknown effects on the transcriptome, we sought to focus on the difference between the control and mutant strains that is elicited by the presence or absence of *daf-42* in the common genetic background of the *daf-2* mutation. Principal component analysis (PCA) showed that while the transcriptome of the *daf-2; daf-42* mutant is similar to that of the *daf-2* control at 52 HAE, the shift of transcriptome made by the *daf-2* control at 60 HAE has not been completely made by the *daf-2; daf-42* mutant (Fig. 5a). Expression of *daf-42* is reduced in the *daf-2; daf-42* double mutant at both 52 and 60 HAE, likely due to nonsense-mediated decay (Fig. 5b, Supplementary Fig. 8a). The difference in transcriptomes at 60 HAE suggests that the absence of *daf-42* hardly affects the transcriptome at 52 HAE, but significantly affects transcriptional changes that occur over the next 8 hours. To determine the status of dauer commitment in *daf-*2 worms at 52 and 60 HAE, we examined genes that can be used to mark dauer commitment within the L2d stage (Shih *et al*. 2019). We found that the expression of *col-183*, which is highly expressed in L2d after dauer commitment, is greatly increased in *daf-2* control larvae at 60 HAE compared to those at 52 HAE (Fig. 5c, Supplementary Fig. 8b). Expression of *col-183* is not altered in *daf-2; daf-42* mutant larvae, implying that dauer commitment is likely to remain intact in the *daf-2; daf-42* mutant and that the action of *daf-42* follows after dauer commitment in L2d.

**Figure 5.**
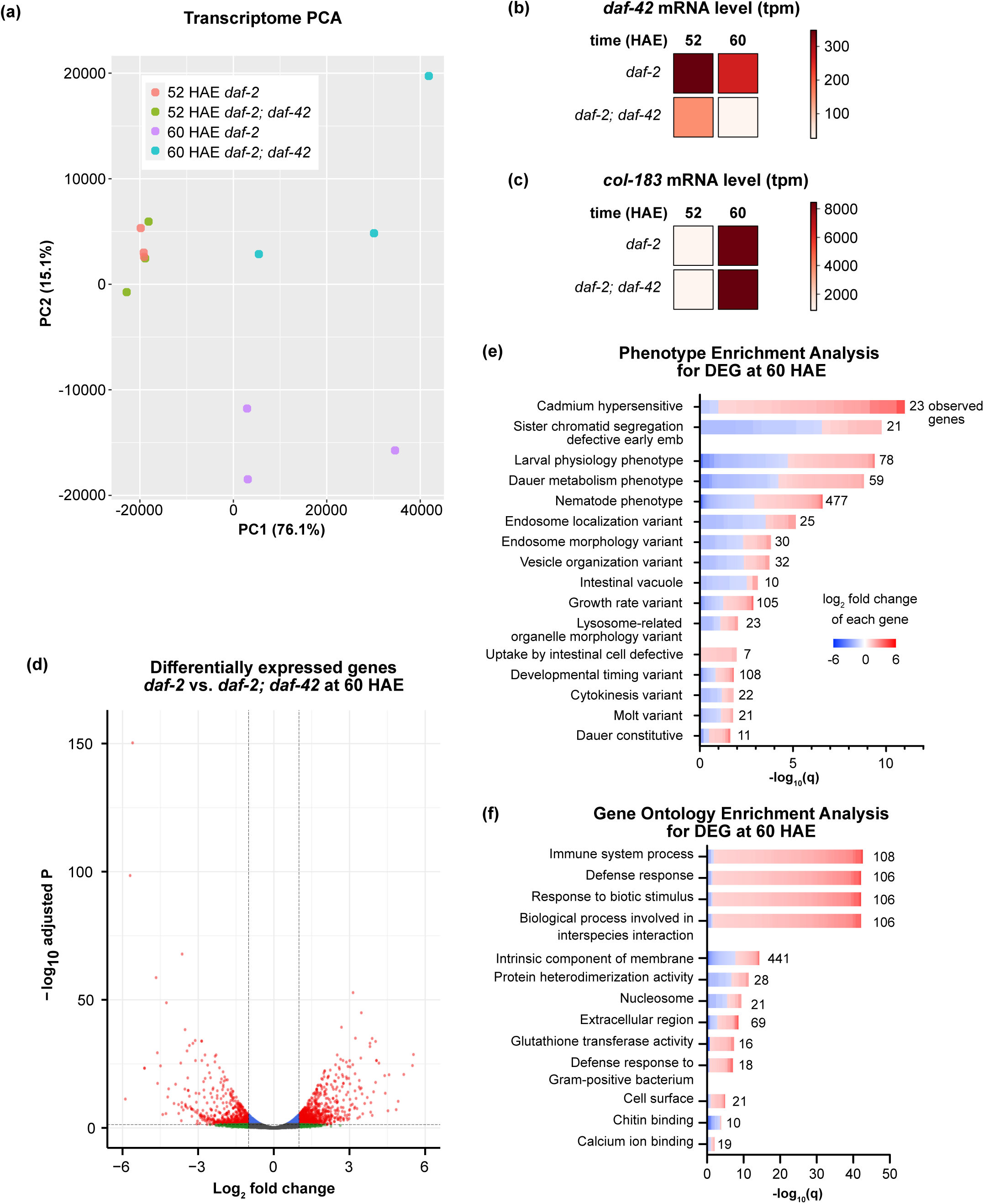
The absence of DAF-42 affects the expression of genes that affect dauer physiology. (a) Principal component analysis (PCA) of the expression profile of *daf-2(e1370)* and *daf- 2(e1370); daf-42(ys54)* strains at 52 HAE and 60 HAE, using transcripts per million (TPM) as the unit of gene expression. (b, c) Average expression levels of *daf-42* (b) and *col-183* (c) at 52 and 60 HAE in *daf-2* control and *daf-2; daf-42* mutant worms. (d, e) Phenotype enrichment analysis (d) and gene ontology enrichment analysis (e) on differentially expressed genes in *daf-2; daf-42* worms compared with *daf-2* worms at 60 HAE. The number of genes detected in each category is on the right side of each bar, and the colored vertical line in each bar represents log_2_-fold difference of each gene within the category from *daf-2; daf-42* mutant compared with the *daf-2* control.

How did the absence of *daf-42* affect the transcriptome? To answer this question, we identified and analyzed differentially expressed genes (DEGs) at 60 HAE. At 60 HAE, 992 genes were significantly upregulated (>2-fold), and 662 genes were significantly downregulated (< 0.5- fold) in the *daf-2; daf-42* mutant compared with the *daf-2* control (Fig. 8d, Supplementary Table 5). Phenotype enrichment analysis of the DEGs between *daf-2* and *daf-2; daf-42* at 60 HAE highlighted that the absence of *daf-*42 significantly alters the expression of genes that affect larval physiology and dauer metabolism (Fig. 5e). While the transcriptome of live L2d worms and dead L2d worms of *daf-2; daf-42* mutant may be different, the gene ontology enrichment analysis indicated upregulation of genes related to the immune system and defense response (Fig. 5f), which likely reflects the consequence of developmental disruption caused by the *daf-42* mutation.

### *daf-42* is an evolutionarily young and essential gene

The results so far indicate that *daf-42* is an essential gene for the development of dauers in *C. elegans*, which is considered to be a stage critical for its survival in the wild. We speculated that *daf-42* would be well-conserved across diverse species and sought to find homologs of *daf-42*.

Contrary to our expectations, a search for DAF-42 homologs on NCBI BlastP revealed no homologs outside nematodes. We searched for DAF-42 homologs in the protein sequences of the 163 nematode species listed in Wormbase Parasite 16 (Parkinson *et al*. 2004; Howe *et al*. 2015; Howe *et al*. 2017; Buchfink *et al*. 2021). Strikingly, highly conserved homologs of DAF-42 m, with bit scores above 100, were found only in *Caenorhabditis* species (Fig. 6a, Supplementary Table 6). This implies that despite its essential role in dauer development, *daf-42* is an evolutionarily young, genus-specific gene.

**Figure 6.**
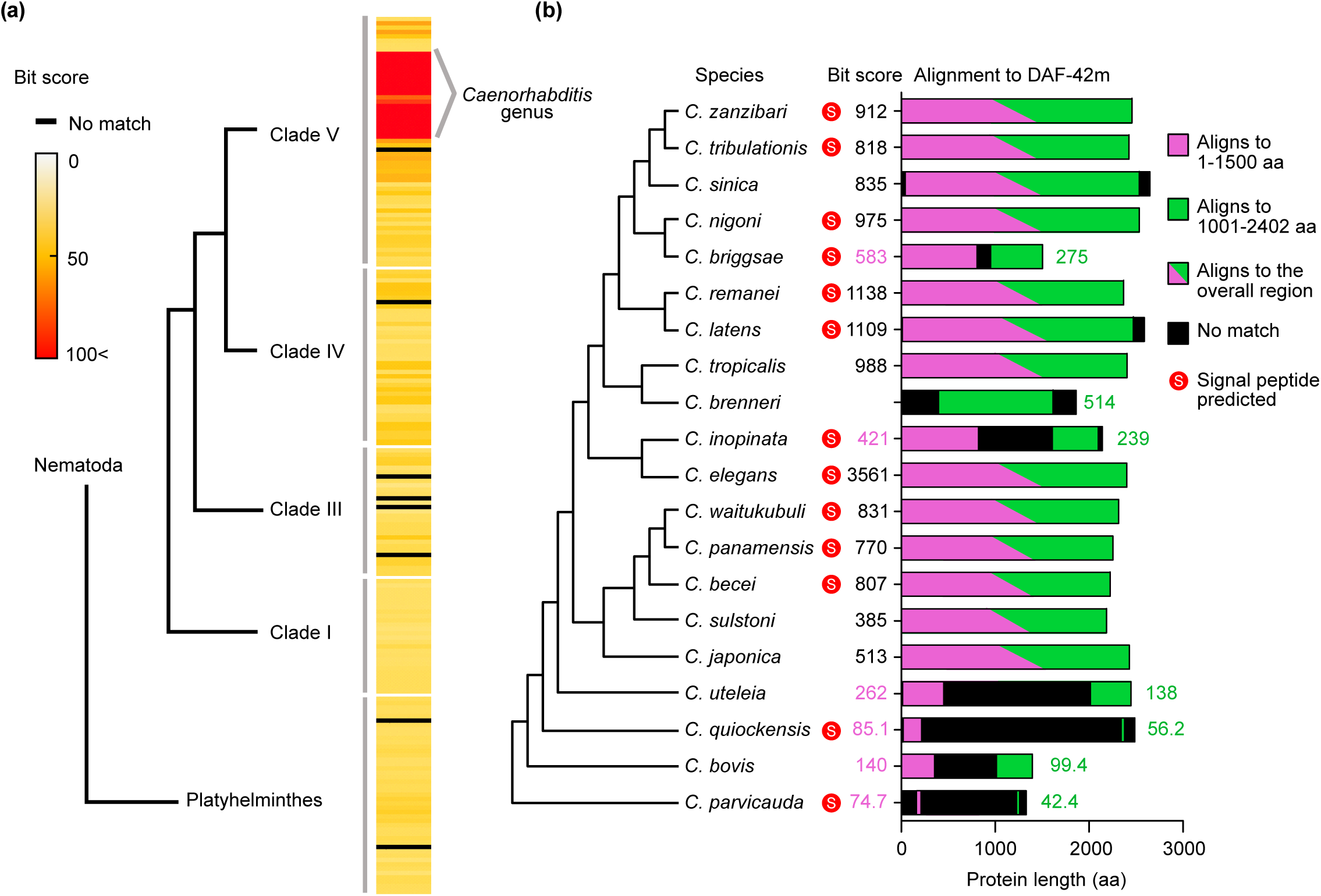
Homologs of DAF-42 in other nematode species. (a) Heat map of bit scores of proteins best-aligned with DAF-42 m from 202 protein database of 163 species in WormBase ParaSite 16. Species within each clade are sorted in alphabetical order of their species name. (b) Diagram of best-matching DAF-42 m homologs in 21 *Caenorhabditis* species. The length of each bar indicates the length of each homolog, and the region that aligns to *C. elegans* DAF-42 is colored. Pink indicates regions that align to 1–1500 amino acid residues of DAF-42 m, and green indicates regions that align with 1,001–2,402 amino acid residues of DAF-42 m. Region that aligns continuously to most length of DAF-42m is marked in both colors. Numbers indicate bit scores of the alignments between DAF-42 m and each homolog; black number indicates the bit score of the alignment between the whole sequences of DAF-42m and the homolog; pink and green numbers indicate the bit score of the alignment of 1–1,500 and 1000–2402 amino acid residues of DAF-42 m with the whole sequence of the homolog, respectively. Homologs predicted to have signal peptides are indicated with red circles with the letter “S”.

Alignment results to *C. elegans* DAF-42 m protein sequence indicated large changes in DAF-42 sequences within *Caenorhabditis* species (Supplementary Table 7). Not only did the four outgroup species have only short conserved regions, but also the *C. brenneri* homolog lacked a region that corresponds to the first half of *C. elegans* DAF-42 m. A comparison of the dot plot between *daf-42* homologs of *C. briggsae* and *C. inopinata* with that of *C. elegans* indicated that the regions in the middle do not align with each other (Supplementary Fig. 9). To confirm this, we divided the DAF-42m sequence into two overlapping halves—regions containing 1–1,500 and 1001–2,402 amino acid residues—and performed protein alignment. We were able to produce similar results (Fig. 6B, Supplementary Table 7). This is particularly interesting because *C. inopinata* phylogenetically closest to *C. elegans* (Kanzaki *et al*. 2018).

Protein dot plots showed that compared to other dauer development-related genes, such as *daf-2* or *daf-7*, *daf-42* underwent more dynamic changes within the genus (Supplemental Fig. 9). In addition, the analysis of protein sequences of *daf-42* homologs in *Caenorhabditis species* for signal peptide also indicated that 6 out of 20 species have lost signal peptide (Fig. 6b). Although all homologs are predicted to contain large proportions of disordered regions, their distributions vary (Supplementary Fig.10, a and b). Despite these changes, amino acid compositions between these homologs remain similar (Supplementary Fig. 10c). These results indicate that while *daf- 42* is an essential gene for dauer development and is conserved only in the *Caenorhabditis* genus, the gene underwent significant changes even within the genus.

## Discussion

In this study, we have newly identified a genus-specific gene that has an essential role in dauer development in *C. elegans*. We found that *daf-42* encodes large, unstructured proteins that are secreted from hypodermal cells in late L2d during the molt into the dauer stage. Notably, although *daf-42* is essential during dauer development—a feature conserved across nematode species—phylogenetic analysis revealed that the *daf-42* is mainly conserved only within the genus *Caenorhabditis*, implicating that *daf-42* is a recently evolved gene that plays an essential role in the survival of the species.

Studies on dauer formation have identified over 30 daf genes that regulate the decision between diapause and reproductive development in late L1 and L2d stages. The environmental and nutritional conditions surrounding the larvae are conveyed by signaling pathways, including *daf-2*/insulin-like, cyclic GMP and *daf-7*/TGF-beta pathways. These pathways converge on steroid hormone signaling mediated by the nuclear hormone receptor *daf-12* and dafachronic acid (Thomas *et al*. 1993; Gottlieb and Ruvkun 1994; Antebi *et al*. 2000; Fielenbach and Antebi 2008). Although *daf* mutants are known to exhibit daf-c and daf-d phenotypes (Fielenbach and Antebi 2008), the *daf-42* mutant displays a completely penetrant lethal phenotype in worms that develop into the dauer stage. As *daf-42* mutation does not affect the developmental decisions of *daf-2*, *daf-7* and *daf-9* mutants (Fig. 1c and Fig. 2, a-c), we hypothesize that *daf-42* belongs to a different category of daf gene that is involved in physiological changes downstream of developmental decisions. The genetic components that mediate the developmental process after the binary decision are unknown, and *daf-42* provides an entry point for further research in this area.

Although the detrimental effect of the *daf-42* mutation in dauer development is clear, the molecular function of DAF-42 protein is unclear. Our results indicate that DAF-42 is a large, disordered protein expressed at late L2d in the seam cells and is secreted towards the surface of the worm. Intrinsically disordered regions are flexible parts of a protein that may provide binding locations and are known to function in various ways, from structural roles to roles in cellular signaling (Peysselon *et al*. 2011; van der Lee *et al*. 2014; Wright and Dyson 2015).

Hypodermal cells, including seam cells, express and secrete various proteins to promote proper cuticle formation and molting. Although the cuticle and proteases are part of the secreted proteins, DAF-42 is expected to be neither because of its enormous size and lack of homology (Frand *et al*. 2005; Chisholm and Xu 2012). A previous study reported that the absence of seam cells hinders body contraction during the dauer stage (Singh and Sulston 1978), which may be relevant in the lack of body contraction and dauer alae in dead L2d *daf-42* mutants. In this context, DAF-42 may be a component of the extracellular matrix or play a signaling role specialized for the dauer stage, that function alone or as a scaffold with its binding partners, which are yet to be identified.

The dauer stage in free-living nematodes is analogous to iL3 in parasitic nematodes in that both are third larval stages with altered physiology and are developmentally arrested until they find a suitable environment to resume development and reproduce (Hotez *et al*. 1993; Hu 2007; Crook 2014). These stages are a crucial part of the life cycle; *C. elegans* is found in the dauer stage in the wild, where resources are often scarce (Frezal and Felix 2015), and iL3 stage plays a special role in host invasion in many parasitic nemaotdes. Studies have reported that *daf-12*, a nuclear receptor that binds dafachronic acid to regulate the decision between dauer and reproductive development programs in *C. elegans* (Antebi *et al*. 2000), is conserved in other nematode species. This function is suspected to be evolutionarily conserved, as the loss of *daf-12* impairs dauer and iL3 development in nematodes in Clade V and IV (Ogawa *et al*. 2009; Dulovic and Streit 2019). *daf-12* homologs in several clade IV and clade III parasitic nematodes were activated by dafachronic acid and other steroid derivatives that affect iL3 development in these species (Wang *et al*. 2009; Ayoade *et al*. 2020; Long *et al*. 2020). By contrast, *daf-42* is different from previous studies in that the gene is conserved only in the genus *Caenorhabditis*, although we may have missed good-matching homolog in other species due to poor quality of protein database. We speculate that nematode species utilize conserved mechanisms involving steroid hormone and *daf-12* in the decision process between diapause and reproductive development, but the mechanisms that mediate physiological changes during the transition into the dauer or iL3 stage may have diverged across various nematode species. A recent study showed that the transcriptome of dauer stage and hypodermis includes more evolutionarily young genes than those of other stages and other tissues in *C. elegans*, respectively (Ma and Zheng 2023). These results indicate that that dauer stage and hypodermis may have been subjected to evolutionary innovation. In this context, *daf-42* may be a part of this innovation that allowed *Caenorhabditis* species to invent the unique features of their dauer stages.

In contrast to the conventional notion that essential genes are evolutionarily old and conserved, studies in the last decade have revealed that young genes also play vital biological roles in diverse species. In *Drosophila*, silencing of young genes that arose in a subgroup of the genus causes critical phenotypes, such as failure in pupal development and sterility (Chen *et al*. 2010; Ding *et al*. 2010; Long *et al*. 2013). In *Caenorhabditis*, telomere-binding protein genes *tebp*, which is a genus-specific gene enriched mostly in the Elegans subgroup, caused sterility when mutated (Dietz *et al*. 2021). Human-specific genes that arose by recent duplication are involved in brain development (Charrier *et al*. 2012). Our findings add another example of young and essential genes.

How did *daf-42* become so important in dauer development? A recent study in *Drosophila* shows that new genes may gain essential function by interacting with other essential genes in the developmental process (Lee *et al*. 2019). Similarly, *daf-42* may be controlled by or bind to more conserved factors to regulate dauer development. Although we are yet to understand the evolutionary origin and role of *daf-42*, we speculate that dauer development in nematodes may be another example of this phenomenon, as *daf-42* is an evolutionarily young gene that plays an essential role in dauer development.

## Data Availability Statement

Strains and plasmids are available upon request. RNA-sequencing data of *daf-2(e1370)* and *daf- 2(e1370); daf-42(ys54)* are submitted to NCBI GEO and are available under accession number GSE230353.

## Acknowledgments

We thank Jiseon Lim for helping with the phylogenetic analysis and the Lee lab members for help and discussion. We also thank Dr. Hee-Jung Choi for helps with the protien structure prediction. Mutant worms were kindly provided by the *Caenorhabditis* Genetics Center (USA).

## Author Contributions

N.K. discovered the “no viable dauer” phenotype. D.S.L., J.K. and J.L. designed the experiments. D.S.L., J.K. and W.K. performed the experiments. S.-H.L. performed electron microscopy. D.S.L., J.K. and D.L. analyzed the data. D.S.L. and J.L. wrote the manuscript.

## References

1. Ahmed, R., Z. Chang, A. E. Younis, C. Langnick, N. Li et al., 2013 Conserved miRNAs are candidate post-transcriptional regulators of developmental arrest in free-living and parasitic nematodes. Genome Biol Evol 5: 1246–1260.

2. Antebi, A., W. H. Yeh, D. Tait, E. M. Hedgecock and D. L. Riddle, 2000 daf-12 encodes a nuclear receptor that regulates the dauer diapause and developmental age in C. elegans. Genes & development 14: 1512–1527.

3. Arribere, J. A., R. T. Bell, B. X. Fu, K. L. Artiles, P. S. Hartman et al., 2014 Efficient marker- free recovery of custom genetic modifications with CRISPR/Cas9 in Caenorhabditis elegans. Genetics 198: 837–846.

4. Ayoade, K. O., F. R. Carranza, W. H. Cho, Z. Wang, S. A. Kliewer et al., 2020 Dafachronic acid and temperature regulate canonical dauer pathways during Nippostrongylus brasiliensis infectious larvae activation. Parasit Vectors 13: 162.

5. Barriere, A., and M. A. Felix, 2005 High local genetic diversity and low outcrossing rate in Caenorhabditis elegans natural populations. Curr Biol 15: 1176–1184.

6. Bray, N. L., H. Pimentel, P. Melsted and L. Pachter, 2016 Near-optimal probabilistic RNA-seq quantification. Nature Biotechnology 34: 525–527.

7. Buchfink, B., K. Reuter and H.-G. Drost, 2021 Sensitive protein alignments at tree-of-life scale using DIAMOND. Nature Methods 18: 366–368.

8. Cassada, R. C., and R. L. Russell, 1975 The dauerlarva, a post-embryonic developmental variant of the nematode Caenorhabditis elegans. Dev Biol 46: 326–342.

9. Charrier, C., K. Joshi, J. Coutinho-Budd, J. E. Kim, N. Lambert et al., 2012 Inhibition of SRGAP2 function by its human-specific paralogs induces neoteny during spine maturation. Cell 149: 923–935.

10. Chen, S., Y. E. Zhang and M. Long, 2010 New genes in Drosophila quickly become essential. Science 330: 1682–1685.

11. Chisholm, A. D., and S. Xu, 2012 The Caenorhabditis elegans epidermis as a model skin. II: differentiation and physiological roles. Wiley Interdiscip Rev Dev Biol 1: 879–902.

12. Consortium, C. e. D. M., 2012 large-scale screening for targeted knockouts in the Caenorhabditis elegans genome. G3 (Bethesda) 2: 1415-1425.

13. Consortium, C. e. S., 1998 Genome sequence of the nematode C. elegans: a platform for investigating biology. Science 282: 2012–2018.

14. Crook, M., 2014 The dauer hypothesis and the evolution of parasitism: 20years on and still going strong. International Journal for Parasitology 44: 1–8.

15. Dietz, S., M. V. Almeida, E. Nischwitz, J. Schreier, N. Viceconte et al., 2021 The double- stranded DNA-binding proteins TEBP-1 and TEBP-2 form a telomeric complex with POT-1. Nat Commun 12: 2668.

16. Ding, Y., L. Zhao, S. Yang, Y. Jiang, Y. Chen et al., 2010 A young Drosophila duplicate gene plays essential roles in spermatogenesis by regulating several Y-linked male fertility genes. PLoS Genet 6: e1001255.

17. Diniz, D. F. A., C. M. R. de Albuquerque, L. O. Oliva, M. A. V. de Melo-Santos and C. F. J. Ayres, 2017 Diapause and quiescence: dormancy mechanisms that contribute to the geographical expansion of mosquitoes and their evolutionary success. Parasit Vectors 10: 310.

18. Doitsidou, M., S. Jarriault and R. J. Poole, 2016 Next-Generation Sequencing-Based Approaches for Mutation Mapping and Identification in Caenorhabditis elegans. Genetics 204: 451–474.

19. Doitsidou, M., R. J. Poole, S. Sarin, H. Bigelow and O. Hobert, 2010 C. elegans Mutant Identification with a One-Step Whole-Genome-Sequencing and SNP Mapping Strategy. PLOS ONE 5: e15435.

20. Dulovic, A., and A. Streit, 2019 RNAi-mediated knockdown of daf-12 in the model parasitic nematode Strongyloides ratti. PLOS Pathogens 15: e1007705.

21. Fielenbach, N., and A. Antebi, 2008 C. elegans dauer formation and the molecular basis of plasticity. Genes Dev 22: 2149–2165.

22. Frand, A. R., S. Russel and G. Ruvkun, 2005 Functional genomic analysis of C. elegans molting. PLoS Biol 3: e312.

23. Frezal, L., and M. A. Felix, 2015 C. elegans outside the Petri dish. Elife 4.

24. Gerisch, B., and A. Antebi, 2004 Hormonal signals produced by DAF-9/cytochrome P450 regulate C. elegans dauer diapause in response to environmental cues. Development 131: 1765–1776.

25. Golden, J. W., and D. L. Riddle, 1984 The Caenorhabditis elegans dauer larva: developmental effects of pheromone, food, and temperature. Dev Biol 102: 368–378.

26. Gottlieb, S., and G. Ruvkun, 1994 daf-2, daf-16 and daf-23: genetically interacting genes controlling Dauer formation in Caenorhabditis elegans. Genetics 137: 107–120.

27. Hand, S. C., D. L. Denlinger, J. E. Podrabsky and R. Roy, 2016 Mechanisms of animal diapause: recent developments from nematodes, crustaceans, insects, and fish. Am J Physiol Regul Integr Comp Physiol 310: R1193–1211.

28. Harris, T. W., V. Arnaboldi, S. Cain, J. Chan, W. J. Chen et al., 2020 WormBase: a modern Model Organism Information Resource. Nucleic acids research 48: D762–D767.

29. Hotez, P., J. Hawdon and G. A. Schad, 1993 Hookworm larval infectivity, arrest and amphiparatenesis: the Caenorhabditis elegans daf-c paradigm. Parasitology Today 9: 23–26.

30. Howe, K. L., B. J. Bolt, S. Cain, J. Chan, W. J. Chen et al., 2015 WormBase 2016: expanding to enable helminth genomic research. Nucleic Acids Research 44: D774–D780.

31. Howe, K. L., B. J. Bolt, M. Shafie, P. Kersey and M. Berriman, 2017 WormBase ParaSite − a comprehensive resource for helminth genomics. Molecular and Biochemical Parasitology 215: 2–10.

32. Hu, P. J., 2007 Dauer. WormBook: 1–19.

33. Jeong, P.-Y., M. Jung, Y.-H. Yim, H. Kim, M. Park et al., 2005 Chemical structure and biological activity of the Caenorhabditis elegans dauer-inducing pheromone. Nature 433: 541–545.

34. Jia, K., P. S. Albert and D. L. Riddle, 2002 DAF-9, a cytochrome P450 regulating C. elegans larval development and adult longevity. Development 129: 221–231.

35. Jumper, J., R. Evans, A. Pritzel, T. Green, M. Figurnov et al., 2021 Highly accurate protein structure prediction with AlphaFold. Nature 596: 583–589.

36. Kanzaki, N., I. J. Tsai, R. Tanaka, V. L. Hunt, D. Liu et al., 2018 Biology and genome of a newly discovered sibling species of Caenorhabditis elegans. Nat Commun 9: 3216.

37. Lazetic, V., and D. S. Fay, 2017 Molting in C. elegans. Worm 6: e1330246.

38. Lee, D., H. Lee, N. Kim, D. S. Lim and J. Lee, 2017a Regulation of a hitchhiking behavior by neuronal insulin and TGF-beta signaling in the nematode Caenorhabditis elegans. Biochem Biophys Res Commun 484: 323–330.

39. Lee, H., M. K. Choi, D. Lee, H. S. Kim, H. Hwang et al., 2011 Nictation, a dispersal behavior of the nematode Caenorhabditis elegans, is regulated by IL2 neurons. Nat Neurosci 15: 107–112.

40. Lee, J. S., P.-Y. Shih, O. N. Schaedel, P. Quintero-Cadena, A. K. Rogers et al., 2017b FMRFamide-like peptides expand the behavioral repertoire of a densely connected nervous system. Proceedings of the National Academy of Sciences 114: E10726–E10735.

41. Lee, Y. C. G., I. M. Ventura, G. R. Rice, D. Y. Chen, S. U. Colmenares et al., 2019 Rapid Evolution of Gained Essential Developmental Functions of a Young Gene via Interactions with Other Essential Genes. Mol Biol Evol 36: 2212–2226.

42. Long, M., N. W. VanKuren, S. Chen and M. D. Vibranovski, 2013 New gene evolution: little did we know. Annu Rev Genet 47: 307–333.

43. Long, T., M. Alberich, F. Andre, C. Menez, R. K. Prichard et al., 2020 The development of the dog heartworm is highly sensitive to sterols which activate the orthologue of the nuclear receptor DAF-12. Sci Rep 10: 11207.

44. Ma, F., and C. Zheng, 2023 Transcriptome age of individual cell types in Caenorhabditis elegans. Proc Natl Acad Sci U S A 120: e2216351120.

45. Madeira, F., M. Pearce, A. R. N. Tivey, P. Basutkar, J. Lee et al., 2022 Search and sequence analysis tools services from EMBL-EBI in 2022. Nucleic Acids Research: gkac240.

46. Mello, C. C., J. M. Kramer, D. Stinchcomb and V. Ambros, 1991 Efficient gene transfer in C.elegans: extrachromosomal maintenance and integration of transforming sequences. The EMBO journal 10: 3959–3970.

47. Mesak, F., A. Tatarenkov and J. C. Avise, 2015 Transcriptomics of diapause in an isogenic self- fertilizing vertebrate. BMC Genomics 16: 989.

48. Necci, M., D. Piovesan, Z. Dosztányi and S. C. E. Tosatto, 2017 MobiDB-lite: fast and highly specific consensus prediction of intrinsic disorder in proteins. Bioinformatics 33: 1402–1404.

49. Ogawa, A., A. Streit, A. Antebi and R. J. Sommer, 2009 A conserved endocrine mechanism controls the formation of dauer and infective larvae in nematodes. Curr Biol 19: 67–71.

50. Okimoto, R., J. L. Macfarlane, D. O. Clary and D. R. Wolstenholme, 1992 The mitochondrial genomes of two nematodes, Caenorhabditis elegans and Ascaris suum. Genetics 130: 471–498.

51. Parkinson, J., M. Mitreva, C. Whitton, M. Thomson, J. Daub et al., 2004 A transcriptomic analysis of the phylum Nematoda. Nature Genetics 36: 1259–1267.

52. Peysselon, F., B. Xue, V. N. Uversky and S. Ricard-Blum, 2011 Intrinsic disorder of the extracellular matrix. Molecular BioSystems 7: 3353–3365.

53. Pimentel, H., N. L. Bray, S. Puente, P. Melsted and L. Pachter, 2017 Differential analysis of RNA-seq incorporating quantification uncertainty. Nature Methods 14: 687–690.

54. Piovesan, D., M. Necci, N. Escobedo, A. M. Monzon, A. Hatos et al., 2021 MobiDB: intrinsically disordered proteins in 2021. Nucleic acids research 49: D361–D367.

55. Ren, P., C. S. Lim, R. Johnsen, P. S. Albert, D. Pilgrim et al., 1996 Control of C. elegans larval development by neuronal expression of a TGF-beta homolog. Science 274: 1389–1391.

56. Riddle, D. L., M. M. Swanson and P. S. Albert, 1981 Interacting genes in nematode dauer larva formation. Nature 290: 668–671.

57. Schaedel, O. N., B. Gerisch, A. Antebi and P. W. Sternberg, 2012 Hormonal signal amplification mediates environmental conditions during development and controls an irreversible commitment to adulthood. PLoS Biol 10: e1001306.

58. Shih, P. Y., J. S. Lee and P. W. Sternberg, 2019 Genetic markers enable the verification and manipulation of the dauer entry decision. Dev Biol 454: 170–180.

59. Singh, R. N., and J. E. Sulston, 1978 Some Observations On Moulting in Caenorhabditis Elegans. Nematologica 24: 63–71.

60. Thein, M. C., G. McCormack, A. D. Winter, I. L. Johnstone, C. B. Shoemaker et al., 2003 Caenorhabditis elegans exoskeleton collagen COL-19: an adult-specific marker for collagen modification and assembly, and the analysis of organismal morphology. Dev Dyn 226: 523–539.

61. Thomas, J. H., D. A. Birnby and J. J. Vowels, 1993 Evidence for parallel processing of sensory information controlling dauer formation in Caenorhabditis elegans. Genetics 134: 1105–1117.

62. Tougeron, K., 2019 Diapause research in insects: historical review and recent work perspectives. Entomologia Experimentalis et Applicata 167: 27–36.

63. van der Lee, R., M. Buljan, B. Lang, R. J. Weatheritt, G. W. Daughdrill et al., 2014 Classification of intrinsically disordered regions and proteins. Chemical reviews 114: 6589–6631.

64. Wang, Z., X. E. Zhou, L. Motola Daniel, X. Gao, K. Suino-Powell et al., 2009 Identification of the nuclear receptor DAF-12 as a therapeutic target in parasitic nematodes. Proceedings of the National Academy of Sciences 106: 9138–9143.

65. Ward, J. D., 2015 Rapid and precise engineering of the Caenorhabditis elegans genome with lethal mutation co-conversion and inactivation of NHEJ repair. Genetics 199: 363–377.

66. Wolkow, C. A., and D. H. Hall, 2011 The Dauer Cuticle, pp. WormAtlas.

67. Wourms, J. P., 1972 Developmental biology of annual fishes. I. Stages in the normal development of Austrofundulus myersi Dahl. J Exp Zool 182: 143–167.

68. Wright, P. E., and H. J. Dyson, 2015 Intrinsically disordered proteins in cellular signalling and regulation. Nature reviews. Molecular cell biology 16: 18–29.

